# GRIMM: GRaph IMputation and Matching for HLA Genotypes

**DOI:** 10.1101/323493

**Authors:** Martin Maiers, Michael Halagan, Loren Gragert, Pradeep Bashyal, Joel Schneider, Polina Lutsker, Yoram Louzoun

**Affiliations:** Center for Blood and Marrow Transplant Research, Minneapolis, MN; National Marrow Donor Program/Be The Match, Minneapolis, MN; Department of Pathology, Tulane University, New Orleans, LA; Department of Mathematics, Bar Ilan University, Ramat Gan, Israel.

## Abstract

**Motivation:** For over 10 years allele-level HLA matching for bone marrow registries has been performed in a probabilistic context. HLA typing technologies provide ambiguous results in that they could not distinguish among all known HLA allele sequences, therefore registries have implemented matching algorithms that provide lists of donor and cord blood units ordered in terms of the likelihood of allele-level matching at specific HLA loci. With the growth of registry sizes, current match algorithm implementations are unable to provide match results in real time.

**Results:** We present here novel computationally-efficient open source implementation of an HLA imputation and match algorithm using a graph database platform. Using graph traversal, our algorithm runtime grows slowly with registry size. This implementation generates results that agree with consensus output on a publicly-available match algorithm crossvalidation dataset.

**Availability:** The Python, Perl and Neo4jJcode is available at https://git.com/nmdp-bioinformatics/grimm

**Supplementary information:** Supplementary data are available at *Bioinformatics* online.

## 1 Introduction

Human leukocyte antigen (HLA) matching between patient and donor is a primary factor determining the success of an allogeneic hematopoietic stem cell (HSC) transplantation (Lee, et al., 2007) (Spellman, et al., 2012). Therefore, the rapid and reliable identification of allele-level HLA matches in unrelated donor registries is of primary importance. This registry search process is performed via a HLA-matching algorithm, which uses population HLA haplotype frequencies to predict which potential matches are most likely to be allele-level matches when additional HLA testing is performed (Dehn, et al., 2016).

Given ambiguous HLA typing data, current match algorithms perform imputation by direct enumeration of all possible haplotype pairs consistent with the typing. The runtime cost of such an algorithm is more than linear with the registry size. With the overall growth in the registry, number of alleles and number of haplotypes in the frequencies set used for imputation, this becomes unfeasible in real time, which is essential for streamlining the donor selection process,

The validate the quality of such algorithms across international registries, the World Marrow Donor Association has developed a formal specification for HLA matching (Bochtler, et al., 2011) and a cross-validation framework for testing matching algorithms (Bochtler, et al., 2016). This framework includes a panel of reference data sets of haplotype frequencies and HLA typing along with collaboratively-generated consensus match results. We here propose novel implementation of imputation and matching based on graph traversal that achieves a cost proportional to the typing ambiguity and total number of potentially-matched donors.

## 2 Methods

The imputation algorithm implementation described here is logically equivalent to previously described methods (Listgarten, et al., 2008; Madbouly, et al., 2014), but is drastically more efficient.

Given an ambiguous HLA typing of a donor and a distribution of population haplotype frequencies, the algorithm calculates the probability of all haplotype pairs consistent with the current typing.

To perform imputation, a graph database with haplotype frequencies for the full set of five HLA loci and also all subsets of locus combinations (partial haplotypes) down to single locus (allele) frequencies is loaded. This graph is constructed with edges connecting each of the partial haplotypes (e.g. the A and C loci) to the top set of full haplotypes, so that partial typing can be directly related to the appropriate full haplotypes (see supplemental methods for more details). This imputation method can tolerate ambiguous and missing data without computing Cartesian products of possible genotypes at missing loci. Our implementation explicitly enumerates phases (possible arrangements of alleles on haplotypes) upfront and uses graph database queries (currently performed using Neo4J - https://neo4j.com/) to deal with allelic ambiguity in the HLA typing data (Fig 1. Upper plot).

**Fig. 1.**
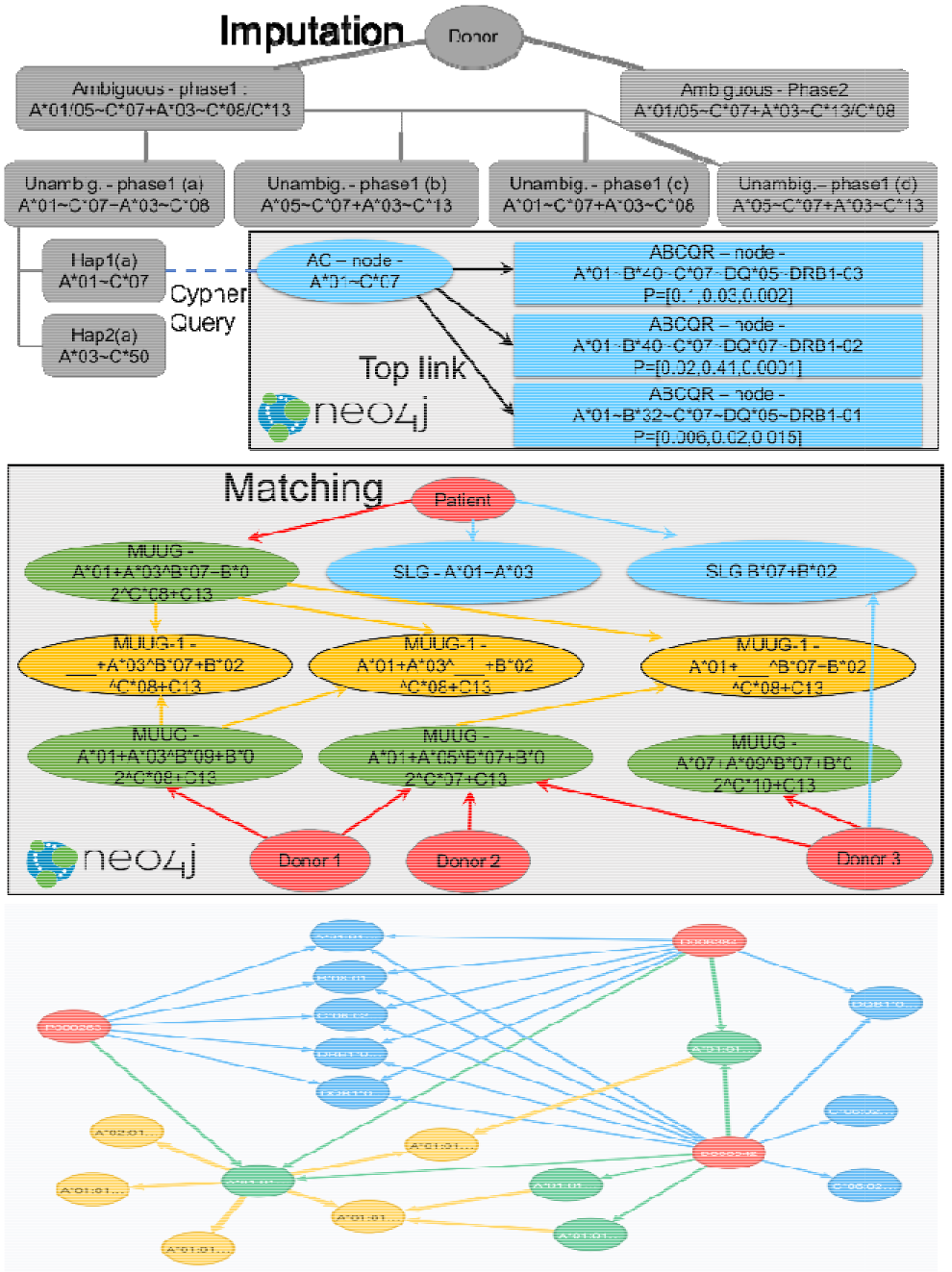
Upper plot schematic model of imputation. Each HLA typing is expanded into all possible ambiguous phases (first row). Each phase is in turn expanded to all possible haplotype pairs. Next a Neo4j query is performed to find all haplotypes in the database consistent with the typing. Lower plot. Schematic model of matching. Each donor is associated with a list of multi-locus unphased genotypes (MUUGs). Finding the donor that can match a patient is simply a graph traversal through the MUUGs (Green circles). For mismatched donors, the traversal must include MUUGs with one or more mismatches (Yellow circles). A parallel query can be performed at the single allele level via singlelocus genotypes (SLGs) (red circles). Central plot – example of actual matching database at high level.

A second graph database is then used to perform matching calculations by representing the results of this imputation process with nodes for each subject (patient, donor) and their associated HLA multi-locus unphased unambiguous genotype (MUUG) and single-locus genotypes (SLG). Edges between each subject and their associated MUUG and SLG nodes are weighted by the likelihood of that subject having the corresponding genotype calculated by imputation (Fig 1 bottom plots). A mathematical description of the method is in the supplementary material.

## 3 Results

We applied this imputation method to the WMDA cross-validation set of simulated patients and donors with the provided haplotype frequencies (Bochtler, et al., 2016). An imputation graph was generated to enumerate probabilities for possible allele-level haplotypes for each subject. Imputation results were then stored in a matching graph that stores MUUGs as nodes with imputed likelihood-weighted edges connecting each subject to their possible allele-level genotypes. We computed match probabilities by running graph traversal Cypher queries against a Neo4J graph database. For example, this query provides the complete list of patient-donor pairs who are potentially full allele matches at 5 loci (10 out of 10) with the probability of match provided as an integer percentage value:

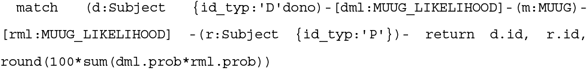

Single allele mismatch searches (e.g. frequency of donors matching in a recipient in nine out of ten alleles) are accomplished by generating MUUG-1 nodes that represent partial genotypes with edges linking them to all corresponding MUUGs. A similar Cypher query provides the integer percentage values for the probability of a single mismatch between a patient and donor at any locus. Single-locus matching is implemented by a similar query for each individual locus based on single-locus genotype (SLG) nodes connected to each subject which were computed during graph construction by summing across all MUUG likelihood edges for that locus.

We generated a set of 10 million match results for each of 1,000 simulated patients compared to 10,000 simulated donors in the WMDA validation dataset. These results were in complete concordance with the cross-registry “consensus” results, except for one donor where we believe the consensus results are incorrect.

The Cypher queries generate results in the order of 100 milliseconds, rather than up to many-minutes from the current match algorithm implementation. This improvement in runtime is largely achieved from storing results from pre-computed imputation sub-problems in the matching graph.

The imputation graph can be generated using any number of population haplotype frequency sets. In the current formalism, very little about the number of HLA loci, resolution, or populations is hard-coded.

Additional categories of HLA matching queries are easy to implement as Cypher queries: haplo-identical matching, multiple-allele-mismatching, or matching only for certain alleles, which could be relevant for targeted T-Cell therapies. This platform has the potential to provide a transparency to matching that is currently lacking and may offer many advantages over current approaches in terms of flexibility in inputs and outputs.

## References

Bochtler, W., et al. A comparative reference study for the validation of HLA Ↄ matching algorithms in the search for allogeneic hematopoietic stem cell donors and cord blood units. Hla 2016;87(6):439–448.

Bochtler, W., et al. World Marrow Donor Association framework for the implementation of HLA matching programs in hematopoietic stem cell donor registries and cord blood banks. Bone marrow transplantation 2011;46(3):338–343.

Dehn, J., et al. HapLogic: a predictive human leukocyte antigen-matching algorithm to enhance rapid identification of the optimal unrelated hematopoietic stem cell sources for transplantation. Biology of Blood and Marrow Transplantation 2016;22(11):2038–2046.

Lee, S.J., et al. High-resolution donor-recipient HLA matching contributes to the success of unrelated donor marrow transplantation. Blood 2007;110(13):4576–4583.

Listgarten, J., et al. Statistical resolution of ambiguous HLA typing data. PLoS computational biology 2008;4(2): e1000016.

Madbouly, A., et al. Validation of statistical imputation of alleleↃlevel multilocus phased genotypes from ambiguous HLA assignments. HLA 2014;84(3):285–292.

Spellman, S.R., et al. A perspective on the selection of unrelated donors and cord blood units for transplantation. Blood 2012;120(2):259–265.

